# The Plastisphere – Marine *fungi* communities in the *plastics age*

**DOI:** 10.1101/2024.09.16.613245

**Authors:** Inga V. Kirstein, Marlis Reich, Yanyan Yang, Maike Timmermann, Antje Wichels, Gunnar Gerdts

**Affiliations:** Alfred-Wegener-Institute Helmholtz Centre for Polar and Marine Research, Biologische Anstalt Helgoland, Helgoland, Germany; University of Bremen, Molecular Ecology, FB2, Leobener Str. 6, 28359 Bremen, Germany

**Author notes:** Correspondence: Inga V. Kirstein.

**Keywords:** Synthetic polymer, Substrate specificity, Community composition, Biofilm, Biodiversity, Multivariate analysis

## Abstract

Fungi play important roles in biofilms, are very versatile in their ecological role, and are considered as plastic degraders. Here we aim to increase the resolution of the fungal members of the Plastisphere, to understand fungal substrate specificities and related potential ecological impacts. Fifteen-month-old fungal Plastisphere communities were investigated on 9 different plastic types and glass in seawater from the North Sea. By integrating scanning electron microscopy (SEM) imaging, ITS-based fingerprinting, and re-evaluated 18S rRNA gene sequence data through a fungal-specific phylogeny-based pipeline, we observed fungal Plastispheres and identified specific characteristics based on morphotypes, phylogeny, and biodiversity across different substrate types. Plastic types selected for specific fungal communities with polyolefine communities indicating significantly higher diversity compared to all other plastic types. Furthermore, specific plastic types may select for specific fungal taxa and their potential hosts, highlighting the complexity of marine biofilm food webs, and related ecological implications.

## 1 Introduction

There are multiple sources and pathways of plastic litter into the ocean but by far, improper disposal of plastics represents the most rapidly growing form of litter entering and accumulating in the oceans (Thiel & Gutow 2005; Andrady 2011). Generally, plastics can be divided into two major groups; plastics with a carbon–carbon backbone and plastics with heteroatoms in the main chain. For example, polyethylene (PE), polypropylene (PP), polystyrene (PS), and polyvinyl chloride (PVC) have a backbone which is solely built of carbon atoms, while polyethylene terephthalate (PET) and polyurethane (PU) feature heteroatoms in the main chain (Gewert *et al*. 2015). The chemical composition (e.g. polyolefines, styrenes, polyesters) and physico-chemical properties of the various plastic types within these two major groups is highly diverse in order to meet the different needs of thousands of products (PlasticsEurope 2019). For example, polyolefines are polymers composed of hydrocarbons of the formula C_n_H_2n_ with a double bond, and are characterised by good chemical resistance and electrical insulation properties. Seven plastic types including high-density polyethylene (HDPE), low-density polyethylene (LDPE), polyvinyl chloride (PVC), polystyrene (PS), polypropylene (PP), polyethylene terephthalate (PET), and polyurethane (PU) make up 81% of the plastics produced in Europe (PlasticsEurope 2019), and are consequently among the most commonly detected plastics in the environment. The estimated 5.25 trillion pieces of plastic currently present in the ocean are a persistent surface acting as substrate for colonisation for a myriad of organisms building up complex biofilms (Dobretsov 2010).

Because plastics are physically and chemically distinct from naturally occurring substrates, they represent a unique surface to the microbial community, and Zettler and colleagues (2013) coined out the term “*Plastisphere*” in order to describe this “new” communities. In the following years, there has been a growing concern about the ecological impact of plastics and it’s Plastisphere on the marine environment. Hence, researchers all over the globe started exploring the microbial Plastisphere at various locations (Oberbeckmann *et al*. 2014; Amaral-Zettler *et al*. 2015; De Tender *et al*. 2015; Bryant *et al*. 2016; Oberbeckmann *et al*. 2016; De Tender *et al*. 2017; Debroas *et al*. 2017; Odobel *et al*. 2021; Latva *et al*. 2022; Yang *et al*. 2022).

Researchers investigating the Plastisphere have mainly focussed on prokaryotic plastic associated communities, pathogens or “*specific*” microorganism/assemblages possibly involved in biological degradation (Zettler *et al*. 2013; Kirstein *et al*. 2016; De Tender *et al*. 2017; Kirstein *et al*. 2018; Oberbeckmann *et al*. 2018; Kirstein *et al*. 2019). Wright *et al*. (2020) demonstrated that Plastisphere communities originating from various habitats could be differentiated between groupings of environmental factors, aspects of study design, and also between plastics when compared with control biofilms. Several microorganisms of diverse environments, including bacteria and fungi, were reported to have a degradative effect on specific plastic types (Restrepo-Flórez *et al*. 2014; Crawford & Quinn 2017; Goudriaan *et al*. 2023). Fungi seem of particular interest in their role as potential plastic degraders in the environment (Krueger *et al*. 2015; Grossart & Rojas-Jimenez 2016), as they are known to be able to degrade several synthetic compounds as e.g., persistent organic pollutants (POPs), and polycyclic aromatic hydrocarbons (PAHs) (Zeghal *et al*. 2021). Furthermore first reports exist on fungal strains degrading plastic (Zeghal *et al*. 2021; Samat *et al*. 2023), originating from various environments like soil, mangroves, lakes and seawater (Zeghal *et al*. 2021). However, in microbial ecology studies, fungi are often only analysed as part of the eukaryotic community, which is why the taxonomic resolution is low. Over the last few years, fungi have come more into the focus of plastisphere researchers and there are about a dozen fungi-specific papers out (Zeghal *et al*. 2021).

Generally, most plastic types are poorly degradable and biological degradation of plastics is known to be slow, as a result plastics remain for years to centuries in marine environments (O’Brine & Thompson 2010). Hence, in order to investigate the Plastisphere reality in the marine environment, long-term exposure studies, as the one by De Tender and colleagues (2017) are needed. However, the studies conducted so far, have either not target to investigate plastic specificities (Davidov *et al*. 2020), or lacked in systematic analysis of distinct plastic types because they focussed on the comparisons of collected secondary marine plastic items/ Plastispheres of unknown exposure time and origin (Lacerda *et al*. 2020), or compared only communities of two plastic types or shapes (De Tender *et al*. 2017; Kettner *et al*. 2017). Kirstein *et al*. (2018) investigated the complete microbial community (pro- and eukaryotes) on seven different types of plastic and a control (glass) of 15-month-old biofilms. The results suggest that fungi may be among the characteristic and discriminatory taxa of the different biofilm communities. Even though the knowledge gained so far provide important first insights into the fungal Plastisphere, we still lack a comprehensive evaluation of the substrate specificity and composition of marine fungal Plastisphere communities. Given that fungi do play important roles in biofilms, are considered as potential plastic degraders, and are very versatile in their ecological function the present study aims to increase the resolution of the fungal members of the microbial plastisphere community as an addition to previously presented findings (Kirstein *et al*. 2018).

In order to investigate the fungal Plastisphere present in the North Sea nine chemically distinct plastic types and glass were exposed in a natural seawater flow-through system experiencing seasonal variation. Following, we investigated 15-month-old fungal communities, using a systematic and statistically robust approach, in order to assess the diversity and potential substrate specificities of fungal Plastisphere communities (Figure 1).

**Figure 1.**
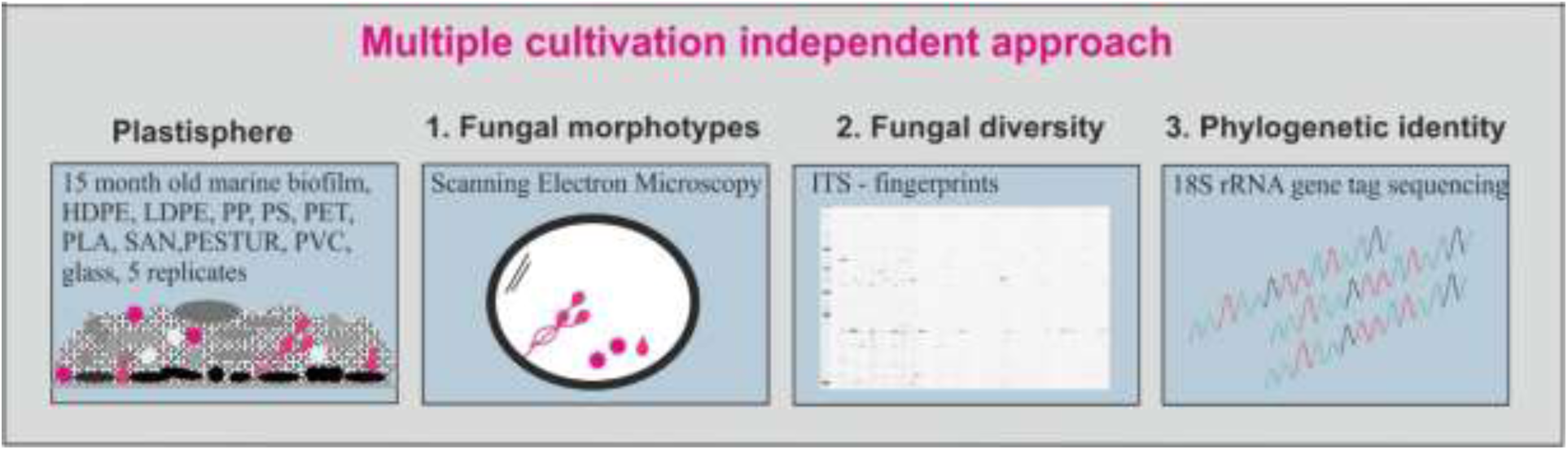
Overview of the study approach to investigate fungal Plastispheres biofilm reality by assessing fungal morphotypes and colonization patterns, diversity and phylogenetic identity.

We used a multiple cultivation independent approach to investigate and characterise fungal Plastisphere communities (Figure 1): 1. Scanning Electron Microscopy (SEM) images of marine plastisphere biofilms were screened for fungal morphotypes and colonization patterns, 2. ITS-based (Internal Transcribed Spacer) fingerprinting of the fungal community was performed to investigate fungal diversity, and 3. 18S rRNA gene sequence data from (Kirstein *et al*. 2018) was re-evaluated with a fungal-specific phylogeny-based pipeline (Banos *et al*. 2020) allowing to resolve phylogenetically also so far undescribed taxa.

## 2 Materials and Methods

### 2.1 Experimental design

Biofilm development was provoked as described previously (Kirstein *et al*. 2018), on 9 chemically distinct plastic types, namely the three polyolefines high-density polyethylene (HDPE), low-density polyethylene (LDPE), and polypropylene (PP), the two styrenes polystyrene (PS) and styrene-acrylonitrite plastics (SAN), polyethylene-tereaphthalate (PET), polylactic acid (PLA), polyurethane prepolymer (PESTUR), polyvinyl chloride (PVC) (Table S1, Figure S1(a), (b)). Biofilms developed on glass slides served as a neutral control. Mimicking plastic present in the North Sea water column, biofilms developed over 15 month in the dark (max. light intensity 0.1033 µmol/m2/s) in a natural seawater flow-through system (Figure S1(a)) located at the island of Helgoland in the German Bight (North Sea, Germany, Latitude 54.18286, and Longitude 7.888838), which is a well-known long-term ecological research site (LTER). Environmental parameters (temperature, salinity, chlorophyll a) of the Seawater flowing through the system, were recorded in the framework of the Helgoland roads long-term data series (Figure S1(d)).

### 2.2 Scanning Electron Microscopy

Scanning electron microscopy was used to visualise the mature biofilms and screen for members of the fungal Plastisphere. Visualisation took place at the Imaging Core Facility of the Julius-Maximillians-University of Würzburg. Sub-samples of each plastic-type and glass of about 0.5 cm^2^ were fixed and processed as described previously (Kirstein *et al*. 2018). A field emission scanning electron microscope (JEOL JSM-7500F) with the in-lens detector (SEI-detector) at 5kV and a working distance of 8 mm was used to visualise the biofilms.

### 2.3 DNA extraction

Biofilm DNA was extracted from each replicate (n = 5) of each substrate (n_substrate_ = 10), as described earlier (Kirstein *et al*. 2018), using a modified protocol from Sapp et al. (2006). Each replicate of each substrate was transferred into 2 mL reaction tubes containing a mixture of 100 μm Zircona/-Silica beads, 700 μL Sodium Chloride –Tris – EDTA (STE) - Buffer was added before mechanically pulped (FastPrep® FP 120, ThermoSavant, Qbiogene, United States). DNA concentrations and purity were determined with a PicoGreen assay (Invitrogen, Waltham, MA) in duplicates using a Tecan Infinite M200 NanoQuant microplate reader (Tecan, Switzerland).

### 2.3 Community profiling using F-ARISA

We exploited the length polymorphism of the ITS1-5.8S-ITS2 region to characterise the fungal community. The ITS region was amplified using the forward primer 2234C (5’-GTTTCCGTAGGTGAACCTGC-3’) and the reverse primer 3126T (5’-ATATGCTTAAGTTCAGCGGGT −3’) (Ranjard *et al*. 2001), the latter labelled with an infrared dye. PCR reactions (10 μL) contained 5 ng template DNA, 2.5 μL Taq Buffer (10x), 500 nM of each primer, 200 μM dNTPs, and 0.5 U Taq DNA polymerase (5 Prime, Hamburg, Germany). Cycling conditions were: 94°C for 3 min, followed by 30 cycles of 94°C for 1 min, 55°C for 30 s and 68°C for 1 min, with a final step at 68°C for 5 min. The PCR products were separated using a LI-COR 4300 DNA Analyser (LI-COR Biosciences, Lincoln, NE, USA) and the analysis of gel images was carried out with the Bionumerics 5.10 software (Applied Maths, Sint-Martens-Latem, Belgium) as previously described by (Krause *et al*. 2012). Bands with intensities lower than 2% of the maximum value of the respective lane and bands smaller than 300 bp were neglected. Incorrect signals defining bands were manually removed. Binning to band classes was performed according to (Kovacs *et al*. 2010). These band classes represent individual taxa and are following referred as Operational Taxonomic Units (OTU).

### 2.4 18S rRNA gene sequencing and taxonomic classification of fungal OTUs

After inspecting the results of the F-ARISA, 18S rRNA gene sequences of all samples obtained (Kirstein *et al*. 2018) were subsequently re-evaluated. In short, sequencing of the hypervariable V4 of the 18S rRNA gene sequence region was performed at LGC Genomics GmbH (Berlin, Germany). Community DNA was amplified using the primer pair Eu565F (5‘-CCAGCASCYGCGGTAATTCC-3⁉) and Eu981R (5‘-ACTTTCGTTCTTGATYRATGA-3’) (Piredda *et al*. 2017). The amplicons were paired-end sequenced 2 x 300 bp on an Illumina MiSeq platform. All sequencing reactions were based upon an Illumina Miseq chemistry following the manufacturer’s instructions.

Quality control of the generated sequences and taxonomic classification was done by using the established sequence pipeline described by (Banos *et al*. 2020). In short, after removal of low-quality sequences, the remaining ones were aligned to the existing alignment of the non-redundant SILVA database SSURef_138.1 (Quast *et al*. 2012) using the SINA algorithm v1.7.2 (Pruesse *et al*. 2012) incorporated in the ARB7 program (Ludwig *et al*. 2004). Only sequences which were classified to fungi based on a 95% sequence similarity threshold were kept and clustered into OTUs based on a 98% similarity threshold using the CD-HIT-EST tool within the CD-Hit software v4.7 (Fu *et al*. 2012). The final classification was done by placing the reference sequences of the OTUs phylogenetically over the Maximum Parsimony algorithm into the fungal phylogenetic 18S rRNA gene sequence reference tree (Yarza *et al*. 2017) using the ARB program. The latest version of the reference tree was used (Priest *et al*. 2021) as it has been enriched by new sequences from the SILVA SSU database and by reference sequences of novel diversity clades described by (Tedersoo *et al*. 2017). Novel diversity clades were defined within the tree if they were formed by at least five sequence reads. Therefore, the lowest possible taxonomic level was transferred and the word “clade new” (e.g. basal fungal clade new) was added. In the case of several new clades formed within a taxon, they were numbered in ascending order. OTUs which were not forming clades, were defined as novel OTU (e.g. Leotiomycetes_unclassfied_OTU_new). In total 492 OTUs (191861 sequences) were identified as fungi, representing 3.36% of all 18S rRNA gene sequences.

Sequence data was deposited in the European Nucleotide Archive (Toribio *et al*. 2017) under the accession number PRJEB22051, using the data brokerage service of the German Federation for Biological Data (Diepenbroek *et al*. 2014), in compliance with the Minimal Information about any (X) Sequence (MIxS) standard (Yilmaz *et al*. 2011).

### 2.5 Statistics and Downstream Data Analysis

For downstream analysis of the 18S rRNA gene sequences, samples with in total less than 50 reads and OTUs with less than 10 reads were rejected from further analysis. A Venn diagram was generated to show differences in OTU and order numbers on plastics *versus* glass using Venny 2.1 (https://csbg.cnb.csic.es/BioinfoGP/venny.html, accessed Feb 2024). The phylogenetic tree of the individual fungal OTUs was annotated with their presence and absence data on the respective substrate using the online tool iTOL version 6.1 (Nayfach *et al*. 2020).

All statistical analyses were carried out with the Primer 7 software including the add-on package PERMANOVA+ (PRIMER-E Ltd, UK) using the presence–absence OTU matrix to avoid overrepresentation of specific taxa in the biofilm (e.g. due to preferential amplification and the presence of uni- and multicellular organisms). Since the taxonomic resolution power of ITS is higher than for the 18S rRNA gene sequence and resolves for a larger number of fungal taxa down to species level (Schoch *et al*. 2012), only the F-ARISA data were used for evaluating differences in alpha diversity. Thus, Shannon diversity (H’ (log2)) was calculated for the individual fungal communities on different plastic types and glass. For beta-diversity analysis and related hypothesis testing of fungal communities, both databases of ITS and 18S rRNA gene sequence, were used as the latter one is more powerful in detecting and classifying undescribed fungal diversity. To visualise patterns in community composition, principal coordinates analysis (PCO) was performed, using the Jaccard index. In order to test for statistically significant variance among the fungal communities attached to different plastics or glass, PERMANOVA with fixed factors and 9999 permutations at a significance level of p<0.05 was performed. To test for within-group dispersion among the substrate groups, separate tests of homogeneity of dispersions (PERMDISP) using 9999 permutations at a significance level of p<0.05 were carried out.

To consider heterogeneity of fungal Plastispheres caused by patchy growth of fungi in the biofilm or due to specificities for a given plastic-type, we created and inspected sub-data sets. Firstly, groups with a known obligate endoparasitic life style (like Rozellomycota) (James & Berbee 2012) were excluded. Next, OTUs present on both plastics and glass were rejected, since they were classified as generalists with no preference. This resulted into a sub-data set with 27 potential plastic “specific” OTUs.

## 3 Results

### 3.1 Plastisphere’s fungal biofilms

After 15 months of exposure to the German Bight’s natural seawater, a dense microbial biofilm was visible on all artificial substrates (Figure S1(c)). Applying scanning electron microscopy a mature multi layered biofilms were covering the substrates evenly (Figure 2) except PLA. PLA biofilms showed locally very thin or an even holey biofilm. Scanning electron microscopic imaging identified different fungal morphotypes on various plastic types (Figure 2).

**Figure 2.**
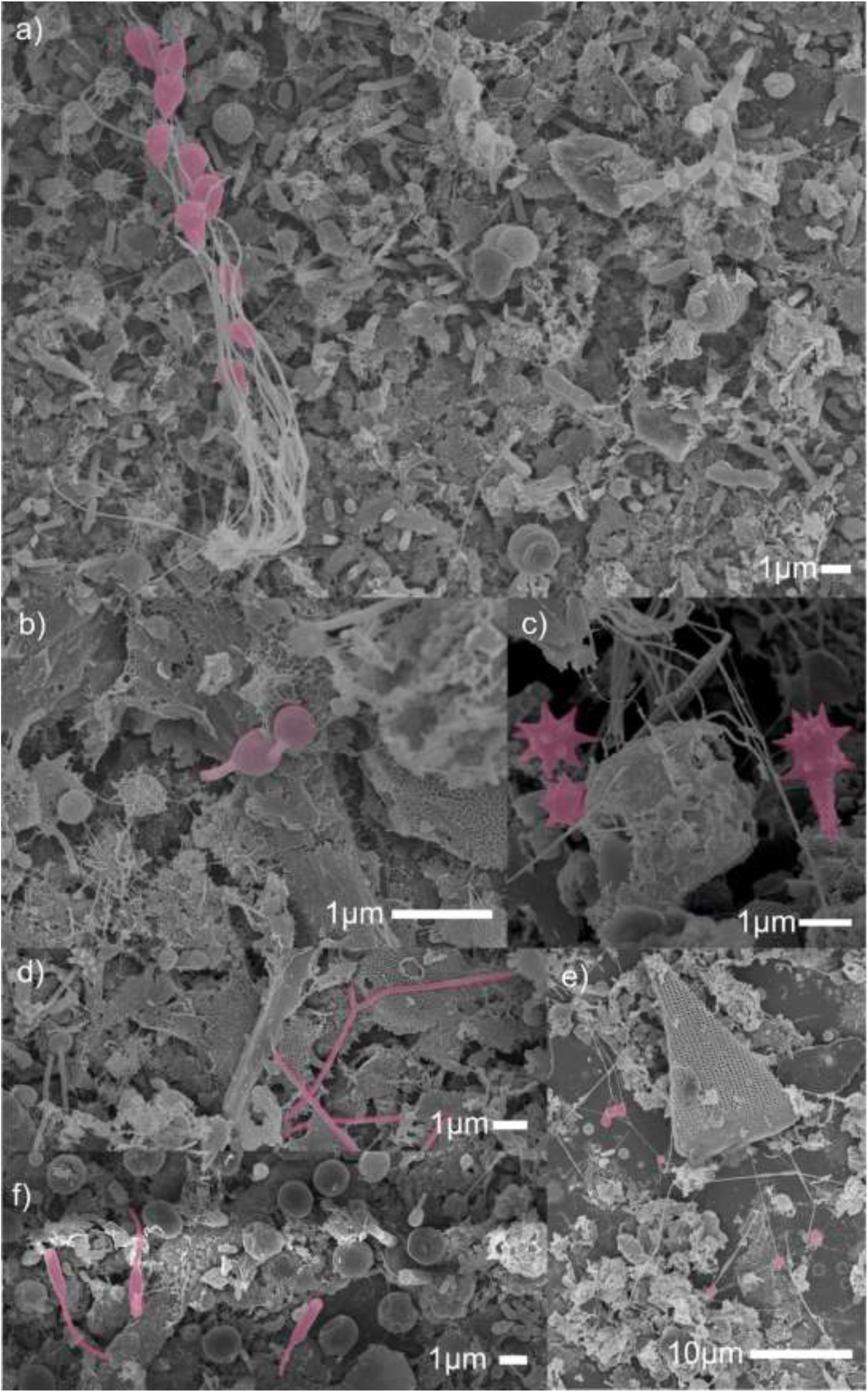
SEM images showing (a) marine fungi as part of a mature biofilm on the surface of PVC, (b) germinating spores on the surface of HDPE, (c) echinulate spores on the surface of LDPE, (d) highly branched, fine hyphal network on the surface of HDPE (e) hyphae with different widths on the surface of glass, lengths and degrees of branching, and (f) fungal spores or conidia on the surface of PLA.

The SEM pictures revealed an extremely high morphological diversity across both prokaryotic and eukaryotic organisms. Fungi were prominent members of the visible microbial community in all biofilms analysed (Figure 2). Different fungal morphotypes were described, such as spores/conidia (Figure 2 (c)), smooth and also ornamented, as well as germinating conidia (Figure 2 (b)). Furthermore, hyphae with different widths, lengths and degrees of branching could be detected (Figure 2 (a)). They were particularly well represented in areas of initial colonisation of the substrate and exhibited a strong 3-dimensionality with their growth forms even in this initial stage (Figure 2 (e). In the mature biofilms, some areas were interwoven with a highly branched, fine hyphal network that entwined other organisms (Figure (d)). Some biofilms were heavily interspersed with fungal spores or conidia (Figure 2 (f)). A preference of the observed morpho-types for a distinct substrate type was not observed and observed fungi appeared patchy within the biofilm.

### 3.2 Alpha-diversity

Analysing the Shannon diversity (H’ (log2) of the different samples revealed that the polyolefine communities had a significantly higher diversity compared to all other plastic types and glass (p<0.05; PERMANOVA, Table S4, Figure 3). Inspecting our results from a chemical perspective, Shannon diversity (H’ (log2)) of F-ARISA fingerprints showed a trend to lower diversity values on plastics with a higher dipole moment in the repeating unit and number of heteroatoms (Figure 3). Furthermore, F-ARISA fingerprints also showed a trend to higher heterogeneity between replicates on plastics with a higher dipole moment in the repeating unit and number of heteroatoms (Figure 3).

**Figure 3.**
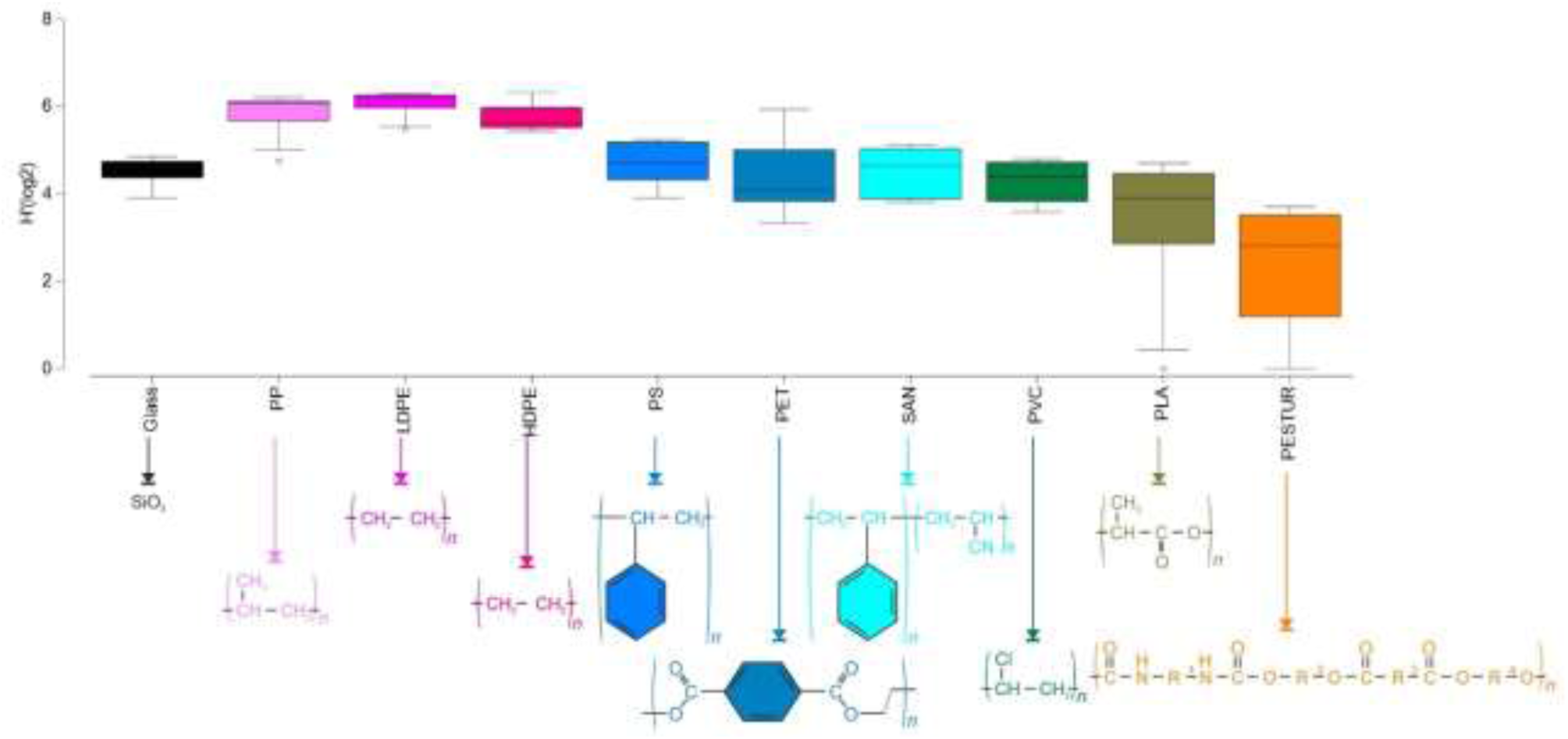
Box-and-whiskers plot of Shannon diversity (H’ (log2)) throughout all plastic types and glass (n = 5). The upper and lower boundaries of the box indicate the 75th and 25th percentiles, respectively. The line within the box marks the median, error bars indicate the 90th and 10th percentiles, and black dots represent outliers. The chemical structure of the substrates is indicated by the colour code, respectively.

### 3.3 Beta-diversity

The taxonomic classification of the fungal communities based on the 18S rRNA gene sequences resulted in a total of 122 OTUs distributed over five different fungal phyla, namely Ascomycota, Basidiomycota, Kickxellomycota, Chytridiomycota, Rozellomycota and two clades of Basal Fungi, which could not be further classified (Figure 4). They grouped into ten classes and nine orders. The undescribed fungal diversity was forming sixteen independent clades all over the phylogenetic tree and on various taxonomic level, from phylum down to order.

**Figure 4.**
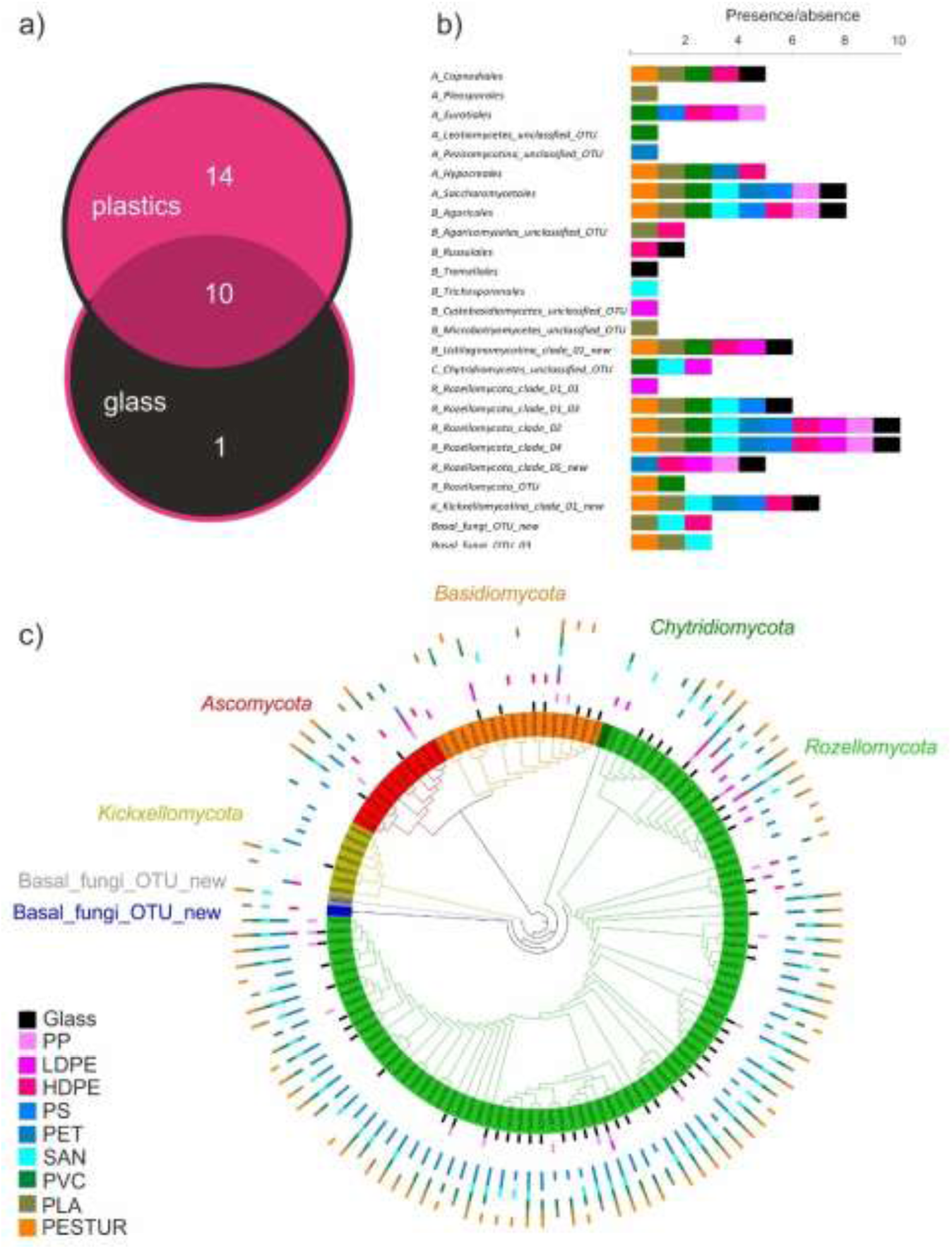
Phylogenetic analysis of fungal biofilms. **(a)** Venn diagram represents fungal orders and in the phylogenetic tree newly formed clades shared and exclusive between the various Plastisphere and glass communities, identified via V4 targeting18S rRNA gene sequence analysis (nOTU_glass_ = 59; nOTU_plastics_ = 118, nOTU_shared_ = 55). **(b)** Fungal orders shown as presence/absence on the distinct plastic types and glass. Substrate types are colour coded, plastics belonging to the group of polyolefins are represented in shades of magenta and pink. A = *Ascomycota*, B = *Basidiomycota*, C = C*hytridiomycota*, R = *Rocellomycota*, K = *Kickxellomycota*. **(c)** Sketch of the phylogenetic tree of the 122 fungal OTUs (18S rRNA gene sequences) detected in this study. The colours of the branches refer to different fungal phyla. The outer rings represent the presence/absence of fungal OTUs on the respective plastic-type or glass by colour.

The phylum Rozellomycota dominated the fungal biofilm communities based on presence/absence data (Figure 4 (c)). However, Rozellomycota was consistently less present on polyolefin surfaces (PP, LDPE, HDPE) than on all other substrates (Figure 4 (c)). We observed members of Ascomycota and Basidiomycota on all plastic types and glass. In contrast, Chytridiomycota was exclusively present on PVC, SAN and LDPE (Figure 4 (c)). When comparing the glass associated fungal communities to pooled fungal Plastisphere communities, 14 fungal orders and new clades defined by the phylogenetic approach were unique to the pooled Plastisphere community while all orders detected on glass were also detected on at least one plastic type with the exception of *Tremellales* (Figure 4 (b)). The ten orders common to pooled plastic communities and/or glass included, besides others, members of *Capnodiales*, *Saccharomycetales*, *Agaricales*, *Russulales*, *Tremellales*, and some clades of *Rozellomycota* (Figure 4 (b), Table S7. Fourteen orders were exclusively detected within plastic communities, including, i.e. *Eurotiales* and *Hypocreales* (Figure 4 (b), Table S7). Out of these 14 orders, seven were exclusively detected on specific plastic types (i.e. *Pleosporales* on PLA, and *Trichosporonales* on SAN, Figure 4 (b)). Only two orders were detected on all plastics and on glass, namely *Rocellomycota*_clade_02 and clade_04 (Figure 4 (b), Table S7). Several OTUs were detected exclusively on only one substrate type, i.e. an unclassified OTU of the sub-phylum *Pezizomycotina* (*Ascomycota*) was only detected on PET (Figure 4 (c)).

### 3.4 Substrate specificity of the fungal Plastisphere communities

The PERMANOVA main tests on F-ARISA fingerprints showed a significant effect (p<0.05; PERMANOVA, Table S2) of the substrate on fungal community structure associated with the various plastics and glass. Following pairwise comparisons showed significant differences between the glass associated biofilm communities and those associated with plastics (p<0.05; pairwise PERMANOVA, Table S3). Significant differences were also observed between polyolefins (PP, LDPE, HDPE) associated fungal communities to all other plastic types but not to each other (Figure 5 (a), (b); Table S3). Furthermore, significant differences were observed between the plastic-pair combinations, PESTUR compared to PS, PET and SAN, and between PS and SAN (Table S3). The PERMANOVA main tests based on 18S rDNA HTS (High Throughput Sequencing) OTUs showed no significant effect (p>0.05; pairwise PERMANOVA, Table S2) of the substrate on fungal community structure associated with the various plastics and glass. However, independent of the chosen approach, F-ARISA and HTS, significant differences were observed between the group of polyolefins (PP, LDPE, HDPE) associated fungal communities compared to all other substrate types (Figure 5 (a), (b); Table S3). With respect to the shade plots generated on the basis of F-ARISA and HTS fungal Plastisphere OTUs, a clear difference in taxonomic resolution was recognisable comparing the two approaches (Figure 5 (c), (d)). Comparing substrate types, more OTUs were observed in the group of polyolefins (PP, LDPE, HDPE) on the basis of F-ARISA compared to all other substrate types (Figure 5 (c)). Vice versa, fewer OTUs were observed in the group of polyolefines on the basis of 18S rDNA HTS OTUs compared to all other substrate types (Figure 5 (d)).

**Figure 5.**
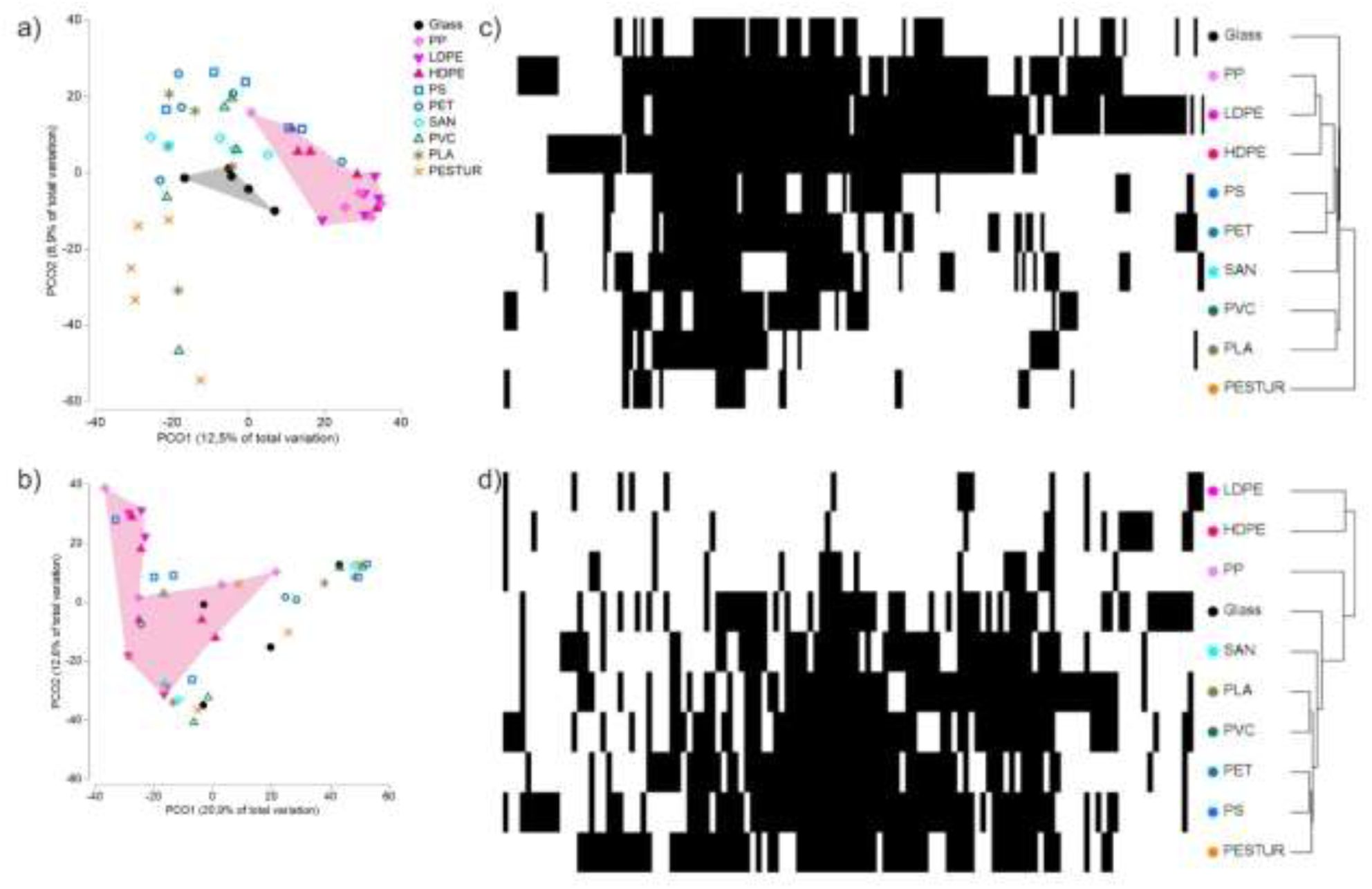
Principle Coordinate Ordination (PCO) relating variation in fungal community composition based on **(a)** F-ARISA OTUs and **(b)** 18S rRNA gene sequences HTS OTUs between different plastic types and glass Plastisphere communities. PCOs represent the similarity of biofilm communities based on the presence/absence of OTUs across samples. Polygons indicate communities present significantly different from all other substrate types (p<0.05; pairwise PERMANOVA, Table S2). Shade Plot of fungal Plastisphere based on **(c)** F-ARISA OTUs and **(d)** 18S rRNA gene sequences HTS OTUs on different plastic types and glass. Shade Plot creation was based on binary data.

In order to unravel potentially plastic-specific taxa, we created a sub-data set containing 27 potential plastic “specific” OTUs. Therefore, *Rozellomycota* which are known to have an obligate endoparasitic life style and generalists present on both plastics and glass (Figure 6) were excluded from the data set. In the tailored data set, nine OTUs belonged to the phylum *Ascomycota* and *Basidiomycota*, respectively. Six OTUs were assigned to *Kickxellomycota*, one OTU was assigned to *Chytridiomycota*, and two OTUs were assigned to Basal Fungi (Figure 6). The PERMANOVA on the generated sub-data set revealed no significant effect (p>0.05; PERMANOVA, Table S6) of the substrate on the potentially plastic specific fungal community structure. Almost 60% (16 out of 27) of the unravelled OTUs were detected exclusively on one specific plastic type. However, out of these 16, 15 OTUs were also detected in a frequency of one of the replicates (n = 5) of a single plastic type. None of the unravelled OTUs was detected on all plastic types. Four OTUs were exclusively detected on PLA belonging to the order *Pleosporales* (*Ascomycota*), an unclassified *Agariomycetes* (*Basidiomycota*), an unclassified *Microbotyomycetes* (*Basidiomycota*), and one OTU belonging to a new clade of *Kickxellomycota*. Furthermore, one OTU belonging to *Cystobasidiomycetes* (*Basidiomycota*) was detected exclusively on LDPE, OTUs belonging to *Eurotiales* (*Ascomycota*), and unclassified *Agariomycetes* (*Basidiomycota*) were solely detected on HDPE, whereas no OTU was exclusively detected on PP. An unclassified OTU assigned to *Pezizomycotina* was detected on PET and *Trichosporonales* (*Basidiomycota*) exclusively on SAN. On PVC, in total, three OTUs were exclusively detected belonging to *Leotiomycetes* and *Saccharomycetales* (both *Ascomycota*) and *Ustilaginomycotina* (*Basidiomycot*a). Two OTUs assigned to *Agaricales* and *Ustilaginomycotina* (both *Basidiomycota*) were exclusively detected on PESTUR. OTUs belonging to a new clade of *Kickxellomycota* were particularly often recognised on PS (5 OTUs out of in total six detected OTUs), suggesting a specificity of the taxa for this plastic-type (Figure 6).

**Figure 6.**
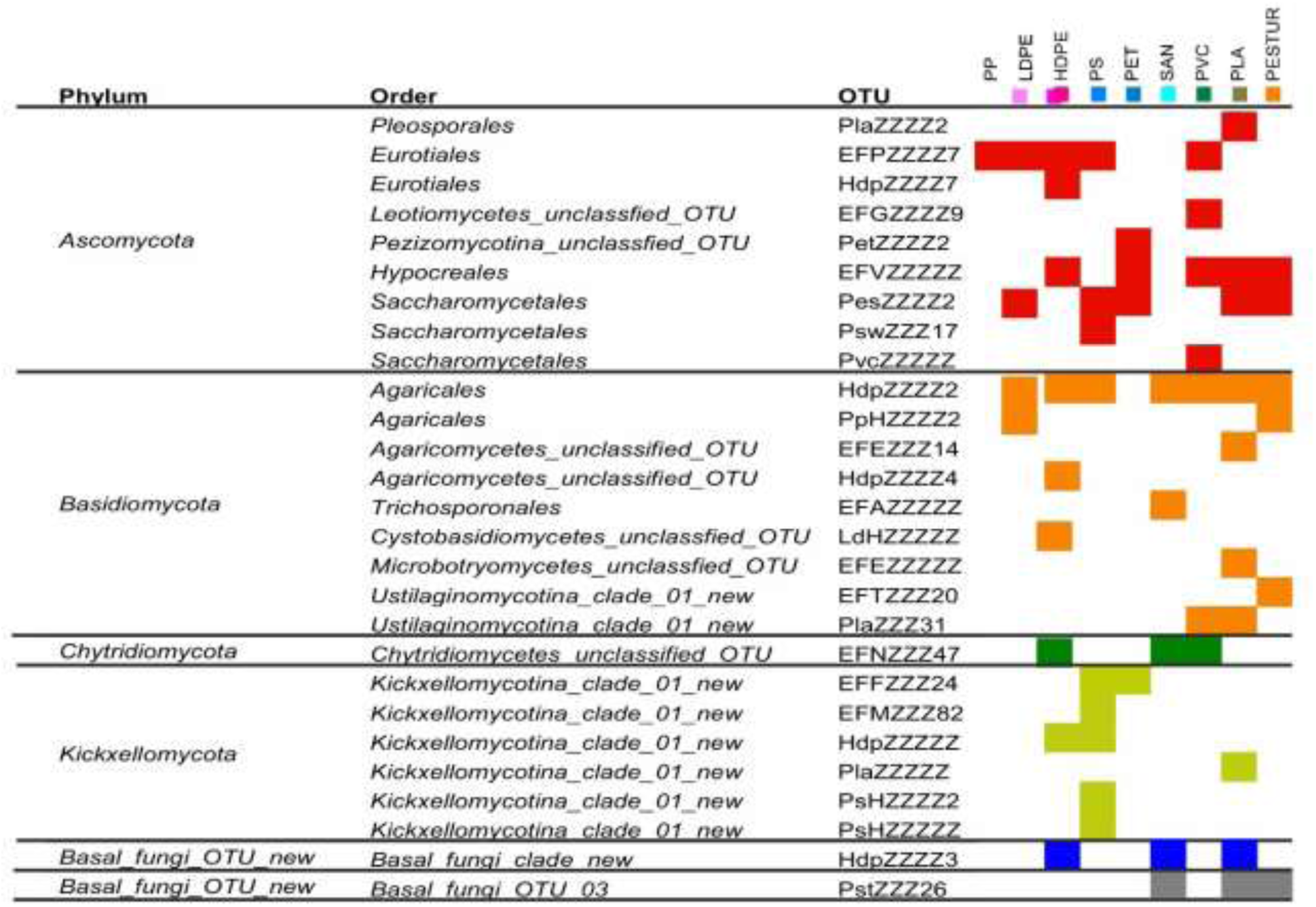
Presence-absence of potentially plastic “specific” fungal taxa on different plastic types. The different colours represent taxa of the same phylum.

## 4 Discussion

### Fungi in the Plastisphere

There are still many open questions about the ecological implications of the ever-growing plastic litter in our oceans, and while several studies addressed the marine plastisphere, its fungal diversity and function remain largely unexplored. In the present study we investigated mature Plastisphere communities observed on plastics in the North Sea using multiple cultivation-independent approaches. Here we will discuss our findings in the context of the following questions. 1. Does plastic select for specific fungal communities? 2. Does plastic serve as accumulation site for specific fungal taxa? 3. What are the potential ecological implications?.

### Does plastic select for specific fungal communities?

In all the used approaches, the structure of fungal biofilm communities was quite heterogenic. Visually, we noticed that fungi appeared in patches within this unique biofilm environment. This confirmed the heterogeneity of fungal biofilm communities both between different samples and across various substrate types, as observed through DNA-based techniques. High variability of fungal Plastisphere communities on PE, also between biological replicates, has been reported previously (De Tender 2017). Both, yeasts and filamentous fungi, can swiftly colonize plastic surfaces, forming biofilms even within minutes (Dusane *et al*. 2008). Surface characteristics like cracks or porosity can expedite colonization rates (Jeyakumar *et al*. 2013). As potential pioneer colonizers, fungi contribute to the formation of diverse biofilms. In cases of active decomposition, fungi may render the carbon source more accessible to other organisms or enrich it by releasing decomposition by-products. Factors like initial colonization and competitive abilities (Heitman *et al*. 2020), including degradation capacities, influence the temporal development of biofilms. This could elucidate the patchiness and high heterogeneity observed across replicates of different plastic types, particularly when filamentous fungi, known for patchy growth, are involved.

Using 18S rDNA HTS no significant differences in community composition between single plastic types or glass were detected. This result is indicating that specificities for particular plastic types are either not present, or hard to detect due to methodological limitations, or are related to the above-mentioned patchy growth of fungi within the biofilm environment. This finding is in consensus with the findings of De Tender and colleagues (De Tender *et al*. 2017). Using ITS2 HTS neither the identification of a fungal core group, nor a clear fungal temporal succession could be observed on PE during their long-term exposure experiment, reflecting the high variability of the fungal communities over time (De Tender *et al*. 2017). However, using F-ARISA we detected significant differences in community composition between several plastic types and glass. Since 18S resolves the basal fungi quite well and ITS primer are known to be biased towards Dikarya, the obtained differences are not necessarily contradicting but rather complementary. Interestingly, all polyolefins were significantly different to all other substrate types, but not to each other. Testing the group of polyolefins revealed significant differences between the group of polyolefins (PP, LDPE, HDPE) associated fungal communities compared to all other substrate types, independent of the chosen approach (F-ARISA and HTS). Comparing PE and PS fungal communities in a short-term exposure experiment Kettner et al. (2017) reported no significant differences (Kettner *et al*. 2017). Even though several studies used the same primers to describe fungal communities previously using F-ARISA (Ranjard *et al*. 2001; Gillevet *et al*. 2009; Krause *et al*. 2012), the primers were found to be homologous to ribosomal sequences from some eukaryotic species that are unrelated to fungi (Ranjard *et al*. 2001; Fechner *et al*. 2010). Consequently, it cannot be completely excluded that other eukaryotic marine organisms were amplified. However, our results clearly indicate that the characteristics or chemical composition of a plastic group / type selected specific fungal communities in the given environment and conditions.

Generally, the composition of biofilm and consequently Plastisphere communities results from various interactions between the biofilms and the surrounding environment, the biofilm age and interaction between diverse organisms within and at the interface of the biofilm, and between biofilms and the substrate they are attached to. De Tender and colleagues carried out a one-year exposure experiment of PE in two different environments, harbour and offshore, in the North Sea, and reported significant effects on fungal communities of both, environment, and exposure time (De Tender *et al*. 2017). Hence, the more uniform conditions present in the natural seawater flow-through system used in the present study, with e.g. less shear forces, may have influenced fungal community composition and subsequent interactions with other organisms.

### Does plastic serve as accumulation site for specific fungal taxa?

The remarkable prevalence of *Rozellomycota* species within the Plastisphere represents a difference to previous studies (Zeghal *et al*. 2021). However, it must be stated, recognising *Rozellomycota* in sequence datasets is still challenging. Initial recognition of this phylum was based on the clustering of sequences from the environment within the phylogenetic tree (Lara *et al*. 2010). To date, there are hardly any sequences that can be linked to culturable isolates. Since the taxonomic classification of fungal taxa in our study was also based on a phylogenetic approach, this methodological choice highlights the differences in recognition and classification compared to other studies.

Presently, the scarcity of cultivable representatives also impedes linking the observed clade formation to a specific taxonomic level. However, estimates suggest a considerable diversity within *Rozellomycota*. These organisms predominantly function as parasites (Wijayawardene *et al*. 2022), exerting significant influence on marine food webs. They act as consumers in both primary and detritus-based food chains, parasitizing primary producers and serving as hyperparasites to primary consumers. The phylum exhibits a wide array of host organisms, infesting phytoplankton, stramenopiles, and other fungi (Gleason *et al*. 2012). This ecological versatility likely contributes to their dominance in the Plastisphere, which might be associated with its mature stage and the diversity of host organisms. However, *Rozellomycota* was consistently less present on polyolefin surfaces than on all other substrates implying that their host may be more abundant on other substrate types.

Zoosporic fungi, including *Rozellomycota*, play pivotal roles in ecosystems by converting host-derived carbon into high-quality long-chain fatty acids (Rasconi *et al*. 2020). Upon release from the host, zoospores become a valuable food source for organisms such as zooplankton. This linkage of different trophic levels in the food chain by fungi is called the mycoloop (Kagami *et al*. 2014). The extent to which this phenomenon applies to zoosporic fungi emerging in the Plastisphere remains uncertain, as data regarding whether zoospores exit the plastisphere to seek new hosts in the pelagic environment is lacking.

In order to unravel potentially plastic-specific taxa, we analysed a sub-data set containing potential plastic “specific” OTUs, excluding the taxa *Rozellomycota* with a known obligate endoparasitic life style and generalists present on both plastics and glass. Several fungal species were previously reported to grow on, change, and/or utilise plastic under specific circumstances, as e.g. *Aspergillus terreus* and *Engyodontium album* (Zeghal *et al*. 2021; Samat *et al*. 2023). However, the biological degradation of plastics is known to be slow and plastics remain, therefore in marine environments for years to centuries (O’Brine & Thompson 2010). The observed plastic “specific” OTUs may also include fungi actively involved in plastic decomposition, although this cannot be conclusively inferred from our dataset. Nonetheless, certain potential plastic utilising representatives of *Ascomycota*, such as *Eurotiales* taxa (Samat *et al*. 2023), were detected within the plastisphere communities studied.

In the fungal plastispheres, especially those associated with PS, additional plastic “specific” unknown taxa of *Kickxellomycota* were detected. This group of fungi is primarily known as intestinal parasites or commensals of marine arthropods, amphipods, and isopods (Heitman *et al*. 2020). While these organism groups are not integral members of the microbial community, it has been observed that some of them colonize floating plastic debris and utilize it as a substrate for attachment or as a habitat (Kar *et al*. 2021). Due to the experimental setup and the size of the plastic films, which provided a stable colonisation surface and were surrounded by natural seawater, colonization of the hosts of *Kickxellomycota* was very likely.

Overall, our results indicate that specific plastic types may select for specific fungal taxa. However, our results also highlight the complexity of unravelling plastic specific fungal taxa in a marine biofilm food web, as the presence of a plastic “specific” fungal taxa could also be related to the presence of a specific host present on the respective plastic type.

### What are the potential ecological implications?

We observed heterogenic diverse fungal communities, unravelled “specific” fungal taxa, and observed significant differences in biodiversity of polyolefine associated fungal communities compared to all other plastic types and glass. Biofilms are generally known to build unique habitats which are often highly diverse. Previously, fungal taxa have been reported to be noticeably more prevalent on plastic (PET) and glass biofilms than in seawater (Oberbeckmann *et al*. 2016) underlining the uniqueness of the biofilm habitat. In the present study, we observed a trend to lower diversity values on plastics with a higher dipole moment in the repeating unit and number of heteroatoms, assuming that the surface polarity and thus wettability might be an important factor for the observed fungal Plastisphere specificities. Li *et al* (Li *et al*. 2021) reported niche-based processes to govern the assembly of plastisphere biofilms, while neutral-based processes dominate the community assembly of the planktonic environment. Hence the question of the ecological implications of the specific fungal Plastispheres associated to distinct plastic types emerges. In order to put our findings into a broader perspective, 250 metric tonnes of plastic is estimated to float in our world oceans (Eriksen *et al*. 2014), serving as a new substrate and surface to colonize for marine organisms. Furthermore, the Plastispheres biomass has been estimated as a fraction of 0.01– 0.2% of the total microbial biomass in the ocean surface (Mincer *et al*. 2019). Plastic serves as a vector for transportation for attached species and thus, associated biomass can be transported from nutrient rich coastal waters to oligotrophic open waters (Mincer *et al*. 2019). This might be especially important as plastic is present in parts of the oceans which is vast and low in biota and organic material. The ingestion of plastics by marine fauna is well documented and its bioavailability dependent on e.g. size, density, abundance, and colour (Wright *et al*. 2013). Hence, the higher number of plastic particles with its associated specific “ecocorona” floating in these areas might change the overall conditions and trigger food web interactions not found at this place so far.

Considering the yearly growing amount of mismanaged plastic litter entering and accumulating in the oceans, the various size fractions, the accumulation and transport of Plastisphere biomass including fungal parasites, and alien species, may have consequences for various ecosystems and for different trophic levels of the food web.

### Conclusion

Our study represents a multiple approach systematic and statistically robust study of mature fungal biofilms associated to distinct plastic types and glass, and therefore enrich our knowledge on the fungal biofilm reality, biodiversity, substrate specificity and taxonomic composition. First and foremost, it has been proofed that mature fungal polyolefine communities differed significantly to all other substrates, independent of the approach used for analysis. Furthermore, we demonstrated that specific plastic types may select for specific fungal taxa or specific hosts, highlighting the complexity of marine biofilm food webs, and related ecological implications. The survival and successful growth of potentially plastic-specific fungi is likely driven by biofilm age, environmental conditions and organism interaction. Optimally, to delineate the effects of season, habitat variation, and substrate specificity on community composition, fungal Plastisphere communities should be analysed at close time intervals over more than one seasonal cycle, and at different locations. To better understand the heterogeneity of fungal biofilm communities, the distribution and location of fungal taxa within the mature biofilm could be assessed by FISH. Further research is needed to assess the ecological implications associated with plastic specific fungal communities.

## Author contribution

I.V. Kirstein together with M. Timmerman performed sampling and DNA extraction. M. Timmerman performed F-ARISA analysis. I.V. Kirstein drafted the manuscript, analysed the re-evaluated 18S rDNA sequences, F-ARISA fingerprints, SEM pictures, and produced figures. M. Reich considerably supported the writing of the manuscript, fungi identification and together with Y. Yang re-evaluated the 18S rDNA sequences. Y. Yang performed the phylogenetic analysis. G. Gerdts and A. Wichels designed the study, and supported the writing of the manuscript. All authors read and contributed to the manuscript.

## Acknowledgements

This study was conducted at the infrastructure Biologische Anstalt Helgoland (Dummermuth *et al*. 2023). We thank all colleagues involved in setting up this experiment and in the day to day running of the station making this work possible. Furthermore, we thank Michaela Meyns for fruitful discussions about plastics chemistry and Hilke Doepke for technical support in the lab. We are grateful for the support in microscopic visualisation of the biofilms by Prof. Dr. Georg Krohne at the University of Würzburg.

