## Supplement for "The Plastisphere – Marine *fungi* communities in the *plastics age*"

*** Correspondence:**

Inga V. Kirstein

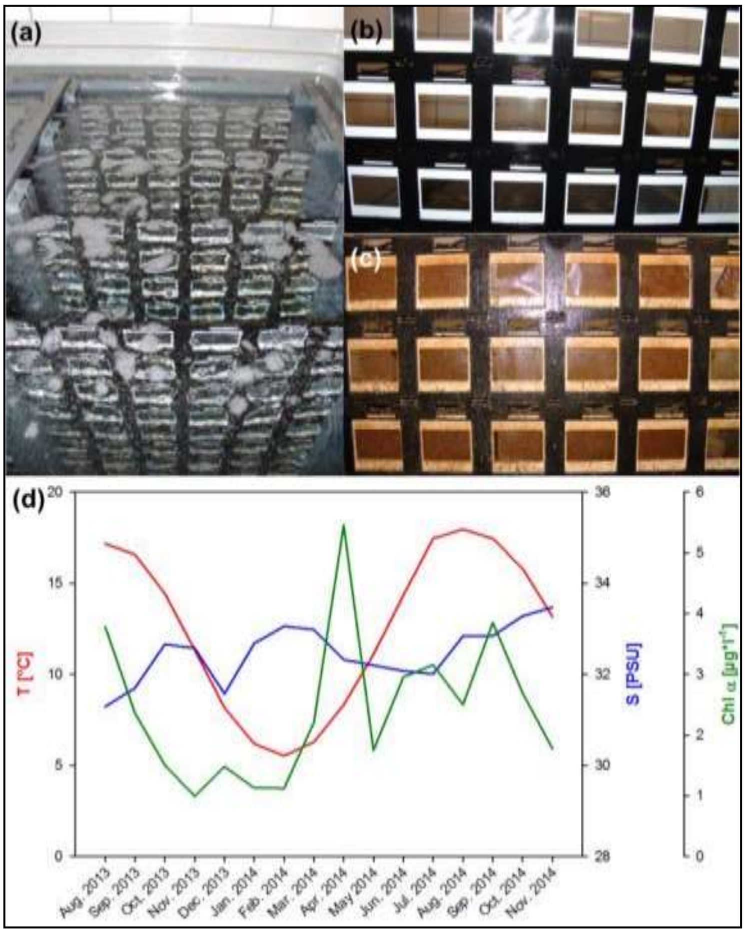


**Figure S1 Flow-through incubation system** for biofilm development on different plastic types and glass mounted in conventional slide-frames. Sea water from Helgoland Roads was directly discharged into the system **(a)** Mounting frames with different plastic foils in the seawater system, **(b)** appearance of the plastic foils at the start in August 2013, **(c)** appearance of the plastic foils in September 2014. **(d)** environmental parameters (monthly means) recorded from 01. August 2013 – 30. November 2014 at Helgoland Roads. T: temperature, S: salinity, Chl *a*: chlorophyll *a*. (adapted from Kirstein et al., 2018).

**Table S1 Sample information** about plastic types used within this study.

| **Polymer** | **Abbreviation** | **Monomer** | **Manufacturer** |
|---|---|---|---|
| Low density polyethylene | LDPE | (C2H4)n | ORBITA-FILM GmbH |
| High density polyethylene | HDPE | (C2H4)n | ORBITA-FILM GmbH |
| Polypropylene | PP | (C3H6)n | ORBITA-FILM GmbH |
| Polystyrene | PS | (C8H8)n | Ergo.fol norflex GmbH |
| Styrene acrylonitrile | SAN | (C8H8)n-(C3H3N)m | Ergo.fol norflex GmbH |
| Polyurethane prepolymer | PESTUR | (C4H4O5)n | Bayer |
| Polylactic acid | PLA | (C3H4O2)n | Folienwerk Wolfen GmbH |
| Polyethylene terephthalate | PET | (C10H8O4)n | Mitsubishi Polyester Film |
| Polyvinyl chloride | PVC | (C2H2Cl)n | Leitz |

**Table S2 PERMANOVA main tests** of F-ARISA ITS fingerprints (n=50) and 18S rRNA V4 tag sequences (n=46) of the fungal biofilm communities on different plastic types and glass based on Jaccard using presence-absence metrics of operational taxonomic units (OTUs). P-values were obtained using type III sums and 9999 permutations under the unrestricted permutation of raw data. d.f.: degrees of freedom, SS: sums of squares; MS: mean squares, perms: number of unique permutations per comparison. [1]Significant results (p *(perm)* < 0.05) are highlighted in bold.

| **18S V4** | | | | | | |
|---|---|---|---|---|---|---|
| **Source of variation** | **d.f.** | **SS** | **MS** | **Pseudo-F** | **p *(perm)*[1]** | **perms** |
| Substrate | 9 | 36728 | 4080,8 | 1,048 | 0,3313 | 9769 |
| Res | 36 | 1,40E+05 | 3894,1 | | | |
| Total | 45 | 1,77E+05 | | | | |
| **ITS** | | | | | | |
| **Source of variation** | **d.f.** | **SS** | **MS** | **Pseudo-F** | **p *(perm)*[1]** | **perms** |
| Substrate | 9 | 47343 | 5260,4 | 1,8144 | **0,0001** | 9630 |
| Res | 40 | 1,16E+05 | 2899,2 | | | |
| Total | 49 | 1,63E+05 | | | | |

**Table S3 PERMANOVA and PERMDISP pair-wise tests** of F-ARISA ITS fingerprints (n=50) of biofilm communities on different plastic types and glass based on Jaccard index, using presence-absence metrics of operational taxonomic units (OTUs). [1]Significant results (p *(perm)* < 0.05) are highlighted in bold.

| | | PERMANOVA | | PERMDISP | |
|---|---|---|---|---|---|
| **Comparison** | | **t *(perm)*** | **p *(perm)*[1]** | **t *(perm)*** | **p *(perm)*[1]** |
| Glass *vs.* | HDPE | 1,4279 | **0,0001** | 1,75 | 0,0701 |
| | LDPE | 1,6024 | **0,0001** | 2,0305 | 0,1068 |
| | PP | 1,4595 | **0,0001** | 1,0054 | 0,4802 |
| | PS | 1,4433 | **0,0001** | 1,4313 | 0,245 |
| | PET | 1,3103 | **0,0001** | 0,58838 | 0,6106 |
| | PLA | 1,3786 | **0,0001** | 0,90589 | 0,5701 |
| | SAN | 1,2739 | **0,0001** | 0,14322 | 0,8949 |
| | PESTUR | 1,4225 | **0,0001** | 2,2885 | 0,0838 |
| | PVC | 1,1992 | **0,0449** | 1,5266 | 0,2716 |
| HDPE *vs.* | LDPE | 1,1211 | 0,1225 | 0,87856 | 0,4481 |
| | PP | 0,98877 | 0,5553 | 0,17913 | 0,8787 |
| | PS | 1,296 | **0,015** | 0,10567 | 0,9198 |
| | PET | 1,2668 | **0,0001** | 2,6756 | **0,0077** |
| | PLA | 1,4314 | **0,0001** | 1,8938 | 0,0945 |
| | SAN | 1,4031 | **0,0001** | 1,8149 | 0,0902 |
| | PESTUR | 1,6813 | **0,0001** | 3,9602 | **0,0096** |
| | PVC | 1,2833 | **0,0083** | 3,4709 | **0,0074** |
| LDPE *vs.* | PP | 0,89684 | 0,7965 | 0,71484 | 0,5767 |
| | PS | 1,6344 | **0,0001** | 0,58228 | 0,7078 |
| | PET | 1,5372 | **0,0001** | 2,7266 | 0,0597 |
| | PLA | 1,6245 | **0,0001** | 2,154 | 0,1051 |
| | SAN | 1,7146 | **0,0001** | 2,0562 | 0,1283 |
| | PESTUR | 1,7628 | **0,0001** | 3,9626 | **0,0071** |
| | PVC | 1,4944 | **0,0001** | 3,4494 | **0,0318** |
| PP vs. | PS | 1,4153 | **0,0075** | 0,21881 | 0,8578 |
| | PET | 1,371 | **0,0001** | 1,539 | 0,2279 |
| | PLA | 1,4474 | **0,0001** | 1,5311 | 0,2515 |
| | SAN | 1,565 | **0,0001** | 0,94265 | 0,417 |
| | PESTUR | 1,6679 | **0,0001** | 2,9224 | **0,0217** |
| | PVC | 1,3721 | **0,0001** | 2,2945 | 0,0949 |
| PS *vs.* | PET | 0,7842 | 0,9349 | 2,0879 | 0,0765 |
| | PLA | 1,1734 | 0,087 | 1,7919 | 0,1918 |
| | SAN | 1,369 | **0,0001** | 1,4041 | 0,2369 |
| | PESTUR | 1,599 | **0,0001** | 3,4628 | **0,0074** |
| | PVC | 1,1684 | 0,0747 | 2,8738 | **0,0237** |
| PET *vs.* | PLA | 0,99107 | 0,5644 | 0,56377 | 0,7647 |
| | SAN | 1,0999 | 0,2524 | 0,79039 | 0,4739 |
| | PESTUR | 1,3975 | **0,0001** | 1,8789 | 0,133 |
| | PVC | 0,89059 | 0,8295 | 1,0359 | 0,4382 |
| PLA *vs.* | SAN | 1,0825 | 0,2384 | 1,0166 | 0,5411 |
| | PESTUR | 1,2082 | 0,0323 | 0,77097 | 0,5373 |
| | PVC | 0,96325 | 0,6385 | 0,1078 | 0,9372 |
| SAN *vs.* | PESTUR | 1,4196 | **0,0001** | 2,5177 | **0,0331** |
| | PVC | 1,0715 | 0,3105 | 1,7634 | 0,1998 |
| PVC *vs.* | PESTUR | 1,1482 | 0,2632 | 0,91456 | 0,4549 |

**Table S4 PERMANOVA main tests** of Shannon diversity (H´(log2) (n=50) F-ARISA fingerprints of biofilm communities on different plastic types and glass based on Euclidean distance. P-values were obtained using type III sums and 9999 permutations under the unrestricted permutation of raw data. d.f.: degrees of freedom, SS: sums of squares; MS: mean squares, perms: number of unique permutations per comparison. [1]Significant results (p *(perm)* < 0.05) are highlighted in bold.

**Shannon**

| **Source of variation** | **d.f.** | **SS** | **MS** | **Pseudo-F** | **p *(perm)*[1]** | **perms** |
|---|---|---|---|---|---|---|
| Substrate | 9 | 59,618 | 6,6243 | 7,6626 | **0,0001** | 9933 |
| Res | 40 | 34,58 | 0,86449 | | | |
| Total | 49 | 94,198 | | | | |

**Table S5 PERMANOVA and PERMDISP pair-wise tests** of Shannon diversity (H´(log2) (n=50) F-ARISA fingerprints of biofilm communities on different plastic types and glass based on Euclidean distance. P-values were obtained using type III sums and 9999 permutations under the unrestricted permutation of raw data. d.f.: degrees of freedom, SS: sums of squares; MS: mean squares, perms: number of unique permutations per comparison. [1]Significant results (p *(perm)* < 0.05) are highlighted in bold.

| | | PERMANOVA | | PERMDISP | |
|---|---|---|---|---|---|
| **Comparison** | | **t *(perm)*** | **p *(perm)*[1]** | **t *(perm)*** | **p *(perm)*[1]** |
| Glass *vs.* | HDPE | 5,3934 | **0,0001** | 0,13868 | 0,9443 |
| | LDPE | 7,01 | **0,0001** | 0,11902 | 0,787 |
| | PP | 4,107 | **0,0001** | 0,90315 | 0,5178 |
| | PS | 0,57668 | 0,5417 | 0,94218 | 0,5048 |
| | PET | 0,57668 | 0,5417 | 1,774 | 0,1279 |
| | PLA | 1,337 | 0,1918 | 2,0274 | 0,0627 |
| | SAN | 0,091107 | 0,9361 | 1,8302 | 0,235 |
| | PESTUR | 3,1496 | **0,0001** | 2,8987 | **0,0166** |
| | PVC | 0,86988 | 0,388 | 1,2354 | 0,434 |
| HDPE *vs.* | LDPE | 1,4692 | 0,1973 | 0,28696 | 0,8088 |
| | PP | 0,20515 | 0,8809 | 0,8568 | 0,7903 |
| | PS | 3,6212 | **0,0072** | 0,90125 | 0,3816 |
| | PET | 2,8873 | **0,0001** | 1,7574 | 0,1752 |
| | PLA | 2,7502 | **0,0001** | 2,0075 | **0,038** |
| | SAN | 3,9788 | **0,0082** | 1,9109 | 0,1053 |
| | PESTUR | 4,9051 | **0,0001** | 2,8993 | **0,0167** |
| | PVC | 5,2378 | **0,0001** | 1,2428 | 0,2316 |
| LDPE *vs.* | PP | 0,82804 | 0,4504 | 1,0315 | 0,4716 |
| | PS | 4,8001 | **0,0001** | 1,0989 | 0,4878 |
| | PET | 3,595 | **0,0001** | 1,8771 | 0,0955 |
| | PLA | 3,1218 | **0,0001** | 2,0693 | 0,0608 |
| | SAN | 5,0666 | **0,0001** | 2,1121 | 0,205 |
| | PESTUR | 5,3751 | **0,0001** | 2,9912 | **0,0162** |
| | PVC | 6,4983 | **0,0001** | 1,4682 | 0,3967 |
| PP vs. | PS | 3,0822 | **0,0084** | 0,069046 | 0,9464 |
| | PET | 2,7419 | **0,032** | 1,0139 | 0,1927 |
| | PLA | 2,7397 | **0,0001** | 1,6572 | 0,4606 |
| | SAN | 3,4462 | **0,0001** | 0,45558 | 0,7586 |
| | PESTUR | 4,7731 | **0,0001** | 2,2187 | 0,082 |
| | PVC | 4,3293 | **0,0001** | 0,0016574 | 1 |
| PS *vs.* | PET | 0,57413 | 0,5846 | 1,1202 | 0,3396 |
| | PLA | 1,4998 | 0,1449 | 1,7079 | 0,3325 |

| | | | | | |
|---|---|---|---|---|---|
| | SAN | 0,54221 | 0,5827 | 0,6111 | 0,5785 |
| | PESTUR | 3,2825 | **0,0077** | 2,3419 | **0,0458** |
| | PVC | 1,2348 | 0,2139 | 0,090585 | 0,9374 |
| PET *vs.* | PLA | 1,0882 | 0,3384 | 1,0585 | 0,6546 |
| | SAN | 0,17276 | 0,8576 | 0,8066 | 0,5049 |
| | PESTUR | 2,5804 | **0,0428** | 1,2194 | 0,2369 |
| | PVC | 0,26094 | 0,7993 | 1,1338 | 0,3222 |
| PLA *vs.* | SAN | 1,2643 | 0,2558 | 1,5458 | 0,484 |
| | PESTUR | 0,9591 | 0,4052 | 0,21117 | 0,9362 |
| | PVC | 1,0336 | 0,3776 | 1,7066 | 0,2718 |
| SAN *vs.* | PESTUR | 2,9677 | **0,0258** | 2,1284 | 0,0542 |
| | PVC | 0,60904 | 0,5281 | 0,63584 | 0,541 |
| PVC *vs.* | PESTUR | 2,7221 | **0,0329** | 2,392 | **0,0225** |

**Table S6 PERMANOVA main tests** of potential plastic specific communities on different plastic types based on Jaccard using presence-absence metrics of operational taxonomic units (OTUs). P-values were obtained using type III sums and 9999 permutations under the unrestricted permutation of raw data. d.f.: degrees of freedom, SS: sums of squares; MS: mean squares, perms: number of unique permutations per comparison.

| **Source of variation** | **d.f.** | **SS** | **MS** | **Pseudo-F** | **p *(perm)*[1]** | **perms** |
|---|---|---|---|---|---|---|
| Substrate | 8 | 16941 | 2117,6 | 0,9453 | 0,6022 | 9844 |
| Res | 33 | 73924 | 2240,1 | | | |
| Total | 41 | 90865 | | | | |

**Table S7 Fungal taxonomic orders based on 18S rRNA** detected in glass, individual and pooled plastic biofilm samples (incubated from August 2013 – November 2014). General sample replication (n=5). Samples with in total less than 50 reads were rejected from analysis resulting in glass (n=4), PLA (n=4), SAN (n=3). Orders are represented by the number of OTUs present on the given substrate; in bold: orders present on both, glass and plastics and sum of detected orders on sample type.

| | **Glass** | **Plastics** | **PP** | **LDPE** | **HDPE** | **PS** | **PET** | **SAN** | **PVC** | **PLA** | **PESTUR** |
|---|---|---|---|---|---|---|---|---|---|---|---|
| A_Capnodiales | **1** | **1** | 0 | 0 | **1** | 0 | 0 | 0 | **1** | **1** | **1** |
| A_Pleosporales | 0 | 1 | 0 | 0 | 0 | 0 | 0 | 0 | 0 | 1 | 0 |
| A_Eurotiales | 0 | 1 | 1 | 1 | 1 | 1 | 0 | 0 | 1 | 0 | 0 |
| A_Leotiomycetes_unclassfied_OTU | 0 | 1 | 0 | 0 | 0 | 0 | 0 | 0 | 1 | 0 | 0 |
| A_Pezizomycotina_unclassfied_OTU | 0 | 1 | 0 | 0 | 0 | 0 | 1 | 0 | 0 | 0 | 0 |
| A_Hypocreales | 0 | 1 | 0 | 0 | 1 | 0 | 1 | 0 | 1 | 1 | 1 |
| A_Saccharomycetales | **1** | **1** | **1** | 0 | 0 | **1** | **1** | **1** | **1** | **1** | **1** |
| B_Agaricales | **1** | **1** | **1** | 0 | **1** | **1** | 0 | **1** | **1** | **1** | **1** |
| B_Agaricomycetes_unclassified_OTU | 0 | 1 | 0 | 0 | 1 | 0 | 0 | 0 | 0 | 1 | 0 |
| B_Russulales | **1** | **1** | 0 | 0 | **1** | 0 | 0 | 0 | 0 | 0 | 0 |
| B_Tremellales | 1 | 0 | 0 | 0 | 0 | 0 | 0 | 0 | 0 | 0 | 0 |
| B_Trichosporonales | 0 | 1 | 0 | 0 | 0 | 0 | 0 | 1 | 0 | 0 | 0 |
| B_Cystobasidiomycetes_unclassfied_OTU | 0 | 1 | 0 | 1 | 0 | 0 | 0 | 0 | 0 | 0 | 0 |
| B_Microbotryomycetes_unclassfied_OTU | 0 | 1 | 0 | 0 | 0 | 0 | 0 | 0 | 0 | 1 | 0 |
| B_Ustilaginomycotina_clade_01_new | **1** | **1** | 0 | **1** | **1** | 0 | 0 | 0 | **1** | **1** | **1** |
| Chytridiomycetes_unclassified_OTU | 0 | 1 | 0 | 1 | 0 | 0 | 0 | 1 | 1 | 0 | 0 |
| Rozellomycota_clade_01_01 | 0 | 1 | 0 | 1 | 0 | 0 | 0 | 0 | 0 | 0 | 0 |
| Rozellomycota_clade_01_03 | **1** | **1** | 0 | 0 | 0 | **1** | 0 | **1** | **1** | **1** | **1** |
| Rozellomycota_clade_02 | **1** | **1** | **1** | **1** | **1** | **1** | **1** | **1** | **1** | **1** | **1** |
| Rozellomycota_clade_04 | **1** | **1** | **1** | **1** | **1** | **1** | **1** | **1** | **1** | **1** | **1** |
| Rozellomycota_clade_05_new | **1** | **1** | **1** | **1** | **1** | 0 | **1** | 0 | 0 | 0 | 0 |
| K_Kickxellomycotina_clade_01_new | **1** | **1** | 0 | 0 | **1** | **1** | **1** | **1** | 0 | **1** | **1** |

| | | | | | | | | | | | |
|---|---|---|---|---|---|---|---|---|---|---|---|
| Basal_fungi_clade_05_new | 0 | 1 | 0 | 0 | 1 | 0 | 0 | 1 | 0 | 1 | 0 |
| Basal_fungi_OTU_03 | 0 | 1 | 0 | 0 | 0 | 0 | 0 | 1 | 0 | 1 | 1 |
| Rozellomycota_OTU | 0 | 1 | 0 | 0 | 0 | 0 | 0 | 0 | 1 | 0 | 1 |
| **Sum** | **11** | **24** | **6** | **8** | **12** | **7** | **7** | **10** | **12** | **14** | **11** |
